# Cebranopadol blocks the escalation of cocaine intake and conditioned reinstatement of cocaine seeking in rats

**DOI:** 10.1101/140566

**Authors:** Giordano de Guglielmo, Alessandra Matzeu, Jenni Kononoff, Julia Mattioni, Rémi Martin-Fardon, Olivier George

## Abstract

Cebranopadol is a novel agonist of nociceptin/orphanin FQ peptide (NOP) and opioid receptors with analgesic properties that is being evaluated in clinical Phase 2 and Phase 3 trials for the treatment of chronic and acute pain. Recent evidence indicates that the combination of opioid and NOP receptor agonism may be a new treatment strategy for cocaine addiction. We sought to extend these findings by examining the effects of cebranopadol on cocaine self-administration (0.5 mg/kg/infusion) and cocaine conditioned reinstatement in rats with extended access to cocaine. Oral administration of cebranopadol (0, 25, and 50 μg/kg) reversed the escalation of cocaine self-administration in rats that were given extended (6 h) access to cocaine, whereas it did not affect the self-administration of sweetened condensed milk (SCM). Cebranopadol induced conditioned place preference but did not affect locomotor activity during the conditioning sessions. Finally, cebranopadol blocked the conditioned reinstatement of cocaine seeking. These results show that oral cebranopadol treatment prevented addiction-like behaviors (i.e., the escalation of intake and reinstatement), suggesting that it may be a novel strategy for the treatment of cocaine use disorder. However, the conditioned place preference that was observed after cebranopadol administration suggests that this compound may have some intrinsic rewarding effects.

## Introduction

Cocaine use disorder is a major health and social problem for which there is no approved medication (Substance Abuse and Mental Health Services Administration, 2010). Most of the pharmacological strategies that have been proposed for the treatment of cocaine addiction have targeted the dopamine system because the acute rewarding effects of cocaine are mediated by the inhibition of dopamine reuptake in the mesocorticolimbic dopamine system (Wise, 1996; Gardner and Ashby, 2000). Unfortunately, these treatment approaches have been ineffective (Grabowski et al., 2000; Haney et al., 2001; Nann-Vernotica et al., 2001; Gorelick et al., 2004). Accumulating evidence indicates that a novel strategy that targets both nociceptin/orphanin FQ peptide (NOP) receptors and μ opioid receptors (MORs) may be useful for the treatment of addictions (Khroyan et al., 2007; Khroyan et al., 2011; Toll, 2013; Kallupi et al., in Press). Buprenorphine is a MOR and NOP receptor partial agonist that has been approved for the treatment of opioid use disorder and has been shown to attenuate the expression of cocaine sensitization in rats (Placenza et al., 2008). Decreases in cocaine self-administration following buprenorphine administration have been reported in both animal models and humans (Mello et al., 1989; Montoya et al., 2004; Sorge and Stewart, 2006b; Wee et al., 2012). These effects have been hypothesized to result from the partial agonist effects of buprenorphine on MORs (Mello et al., 1993), which confers buprenorphine lower abuse liability compared with morphine and is reflected by its classification as a Schedule III compound by the United States Drug Enforcement Administration. However, buprenorphine is also a partial agonist at the NOP receptor (Spagnolo et al., 2008; Khroyan et al., 2009). Its effects on NOP receptors may explain some of its therapeutic effects and lower addiction potential. For example, NOP receptor agonists have been reported to increase the analgesic effects of MOR agonists (Courteix et al. 2004; Cremeans et al. 2012), decrease dopamine release from neurons in brain areas that are associated with the reward pathway (Ikemoto and Panksepp, 1999; Toll, 2013), decrease morphine-induced conditioned place preference (Ciccocioppo et al., 2000), and attenuate the rewarding effects of opioids in rats (Murphy et al., 1999; Rutten et al., 2010). Interestingly, recent results showed that buprenorphine requires concomitant activation of NOP and MOP receptors to reduce cocaine consumption in rats (Kallupi et al., in Press). This evidence suggests that compounds with high efficacy at MORs and NOP receptors may be useful for the treatment of drug addiction. Importantly, however, a specific NOP receptor agonist failed to prevent cocaine-induced conditioned place preference (CPP), the extinction of cocaine-induced CPP, and the stress- and cocaine-induced reinstatement of cocaine seeking (Sartor et al., 2016), suggesting that specific NOP receptor agonists may not be a relevant strategy for the treatment of cocaine use disorder.

The present study tested the hypothesis that oral administration of cebranopadol (trans-6′-fluoro-4’,9’-dihydro-*N*,*N*-dimethyl-4-phenyl-spiro[cyclohexane-1,1’(3’*H*)-pyrano[3,4-*b*]indol]-4-amine), a small molecule that targets opioid receptors, has preclinical efficacy in reducing addiction-like behavior in an animal model of cocaine use disorder. Cebranopadol is a novel opioid receptor agonist with analgesic properties, full agonist activity at NOP receptors and MORs, and partial agonist activity at k opioid receptors and δ opioid receptors (Linz et al., 2014; Schunk et al., 2014). Cebranopadol has been shown to exert potent and effective analgesic and antihypersensitive effects in several rat models of acute and chronic pain (e.g., tail-flick test, rheumatoid arthritis, bone cancer, spinal nerve ligation, and diabetic neuropathy). It has a long duration of action (> 9 h after oral administration of 55 μg/kg in the rat tail-flick test). Unlike morphine, cebranopadol does not disrupt motor coordination or respiration at doses that are within or exceed the analgesic dose range (Linz et al., 2014; Schunk et al., 2014; Lambert et al., 2015). Given these unique pharmacological characteristics, we tested the efficacy of cebranopadol in inhibiting the escalation of cocaine intake and preventing cocaine-seeking behavior in rats with a history of extended access to cocaine self-administration. We also evaluated the potential rewarding effects of cebranopadol by measuring cebranopadol-induced conditioned place preference. Finally, we evaluated the behavioral specificity of cebranopadol by comparing its effects on cocaine-directed behavior with behavior that was maintained by a highly palatable conventional reinforcer (sweetened condensed milk [SCM]).

## Materials and Methods

### Animals

Male Wistar rats (Charles River, Wilmington, MA, USA; *n* = 45) were housed two per cage on a reverse 12 h/12 h light/dark cycle (lights off at 8:00 AM) in a temperature (20–22°C) and humidity (45–55%) controlled vivarium with *ad libitum* access to tap water and food pellets (PJ Noyes Company, Lancaster, NH, USA). All of the procedures were conducted in strict adherence to the National Institutes of Health *Guide for the Care and Use of Laboratory Animals* and were approved by the Institutional Animal Care and Use Committee of The Scripps Research Institute. At the time of testing, the rats’ body weight ranged between 350 and 400 g.

### Intravenous catheterization

The animals were anesthetized by inhalation of a mixture of isoflurane, and intravenous catheters were aseptically inserted in the right jugular vein using a modified version of a procedure that was described previously (Caine and Koob, 1993; de Guglielmo et al., 2013). The vein was punctured with a 22-gauge needle, and the tubing was inserted and secured inside the vein by tying the vein with suture thread. The catheter assembly consisted of a 18 cm length of Micro-Renathane tubing (0.023 inch inner diameter, 0.037 inch outer diameter; Braintree Scientific, Braintree, MA, USA) that was attached to a guide cannula (Plastics One, Roanoke, VA, USA). The guide cannula was bent at a near right angle, embedded in dental acrylic, and anchored with a mesh (2 Å thick, 2 cm square). The catheter exited through a small incision on the back, and the base was sealed with a small plastic cap and metal cover cap. This design helped to keep the catheter base sterile and protected. The catheters were flushed daily with heparinized saline (10 U/ml of heparin sodium; American Pharmaceutical Partners, Schaumburg, IL, USA) in 0.9% bacteriostatic sodium chloride (Hospira, Lake Forest, IL, USA) that contained 20 mg/0.2 ml of the antibiotic Timetin (GlaxoSmithKline).

### Drugs

Cebranopadol (Med Chem Express, Monmouth Junction, NJ, USA) was dissolved in vehicle that consisted of 5% dimethylsulfoxide, 5% Emulphor, and 90% distilled water. The solution was vortexed before filling a 1-ml syringe for oral injection. Cebranopadol was injected orally by gavage at doses of 0.0, 25, and 50 μg/kg. Cocaine HCl (National Institute on Drug Abuse, Bethesda, MD, USA) was dissolved in 0.9% saline (Hospira, Lake Forest, IL, USA) at a dose of 0.5 mg/kg/infusion and self-administered intravenously. Sweetened condensed milk (Nestlé, Solon, OH, USA) was diluted 2:1 (v/v) in water.

### Operant training

Self-administration was performed in operant conditioning chambers (Med Associates, St. Albans, VT, USA) that were enclosed in lit, sound-attenuating, ventilated environmental cubicles. The front door and back wall of the chambers were constructed of transparent plastic, and the other walls were opaque metal. Each chamber was equipped with two retractable levers that were located on the front panel. Cocaine was delivered through plastic catheter tubing that was connected to an infusion pump. The infusion pump was activated by responses on the right (active) lever. Responses on the left (inactive) lever were recorded but did not have any scheduled consequences. Activation of the pump resulted in the delivery of 0.1 ml of the fluid. A computer controlled fluid delivery and behavioral data recording.

For the studies of SCM self-administration, each chamber was equipped with a reservoir that was positioned 4 cm above the grid floor in the center of the front panel of the chamber. Two retractable levers were each located 3 cm away from the SCM receptacle (one lever on the right and one on the left). A pump was activated by responses on the right (active) lever, and responses on the left (inactive) lever were recorded but did have scheduled consequences. Active lever presses resulted in the delivery of 0.1 ml of SCM. A computer controlled fluid delivery and recorded the total number of rewards and number of right and left lever presses.

### Effect of cebranopadol on the escalation of cocaine self-administration

Rats (*n* = 14) were trained to self-administer cocaine under a fixed-ratio 1 (FR1) schedule of reinforcement in daily 6-h sessions. Each active lever press resulted in the delivery of one cocaine dose (0.5 mg/kg/0.1 ml infusion). A 20-s timeout (TO) period followed each cocaine infusion. During the timeout period, responses on the active lever did not have scheduled consequences. This TO period occurred concurrently with illumination of a cue light that was located above the active lever to signal delivery of the positive reinforcement. The rats were trained to self-administer cocaine in 15 sessions (5 days/week) until a stable baseline of reinforcement was achieved (< 10% variation over the last three sessions). A within-subjects Latin-square design was used for the drug treatments. The rats were orally injected with cebranopadol (0, 25, and 50 μg/kg) 30 min before beginning the sessions. Oral administration was performed by gavage using a 19-gauge needle and 8 cm of Tygon tubing (0.030 inch inner diameter, 0.090 inch outer diameter). The animals were subjected to cocaine self-administration at 2 day intervals between drug tests.

### Effect of cebranopadol on sweetened condensed milk self-administration

Rats (*n* = 12) were trained to self-administer SCM under an FR1 schedule of reinforcement for 6 h per day to match cocaine self-administration. Following each SCM reward delivery, a 20-s TO period occurred, during which responses on the active lever had no scheduled consequences. This TO period occurred concurrently with illumination of a cue light that was located above the active lever to signal delivery of the positive reinforcement. The rats were trained to self-administer SCM for several days until a stable baseline of reinforcement was achieved (< 10% variation over the last three sessions). When the stable baseline was reached, the rats orally received cebranopadol (0, 25, and 50 μg/kg) 30 min before beginning the next session. The animals were subjected to SCM self-administration at 2 day intervals between drug tests.

### Cebranopadol-induced conditioned place preference and locomotor activity

Cebranopadol-induced CPP was evaluated using a biased, counterbalanced CPP procedure. Naive rats (*n* = 9) were handled and habituated to oral administration for 1 week before beginning the study. The experiment consisted of three 30-min phases: pretest (one session), conditioning (eights sessions), and preference test (one session). The rats were placed in a dim (40 lux) room 30 min before starting the tests. A two-chambered (38 cm × 32 cm × 32 cm) place conditioning apparatus was used, with both visual cues on the walls (stripes or dots for compartments A and B, respectively) and tactile cues on the floor (smooth or rough for compartments A and B, respectively). On day 1 (pretest), naive rats were placed between the two chambers and allowed to freely explore both chambers for 30 min. Individual bias toward either compartment A or B was observed, and the animals were assigned to place conditioning subgroups according to their least preferred compartment; therefore, biased assignment was used. Conditioning was performed within-subjects. Each rat received cebranopadol (25 μg/kg, p.o.) and vehicle on alternating days in a counterbalanced design 30 min prior to being placed in the conditioning chamber. The preference test was performed 24 h after the last conditioning session. Thirty minutes after vehicle administration, the rats were placed in the non-conditioned side of the apparatus with free access to both chambers. The time spent in the different chambers was recorded. Locomotor activity was recorded in each phase (pretest, conditioning, and preference test) using a video camera that was connected to the ANY-maze Video Tracking System 5.11.

### Effect of cebranopadol on conditioned reinstatement of cocaine-seeking behavior

#### Cocaine self-administration training

Rats (*n* = 10) were surgically prepared with indwelling micro-urethane catheters that were inserted in the right jugular vein. Following 7 days of postsurgical recovery, the rats began self-administration training. The rats were trained to self-administer cocaine (0.5 mg/kg/0.1 ml infusion, i.v.) for 6 h/day on an FR1 schedule in the presence of a contextual/discriminative stimulus (S^D^). Each session was initiated by extending two retractable levers into the operant conditioning chamber. Constant 70 dB white noise served as a discriminative stimulus that signaled availability of the reinforcer throughout the session. Responses on the right, active lever were reinforced with a dose of cocaine, followed by a 20-s TO period that was signaled by illumination of a cue light above the active lever. During this TO period, the lever remained inactive to prevent accidental overdosing with cocaine. Responses on the left, inactive lever had no scheduled consequences.

#### Extinction

Following completion of the training procedure (21 sessions), the extinction phase began. The rats were subjected to daily 2 h extinction sessions, in which responses on the previously active lever had no programmed consequences (i.e., no cocaine delivery and no cue presentation). This phase lasted until responding was extinguished (< 10 responses per session for 3 consecutive days).

#### Conditioned reinstatement

Twenty-four hours after the last extinction session, all of the rats were presented with neutral stimuli (S^N^) in a 2 h session to control for the specificity of the S^D^ to reinstate extinguished cocaine-seeking behavior. The neutral stimuli (S^N^) that signaled the non-availability of the reinforcer consisted of illumination of a 2.8 W house light that was located on top of the chamber’s front panel for the entire duration of the session (2 h). Responses at the right active lever were followed by the presentation of a 20 s intermittent tone, during which the lever remained inactive.

Two days later, the rats were presented with the S^D^. To evaluate the effect of cebranopadol on the conditioned reinstatement of cocaine seeking, the rats were treated with cebranopadol (0, 25, and 50 μg/kg) in a counterbalanced Latin-square design 30 min before the reinstatement test. The reinstatement test lasted 2 h under S^D^ conditions, except that cocaine was unavailable. Cebranopadol was administered only in the S^D^ conditions, with a 2-day interval between tests.

### Statistical analysis

The effects of cebranopadol on cocaine self-administration, SCM self-administration, and conditioned reinstatement and differences in responding during the extinction, S^N^, and S^D^ sessions were analyzed using one-way within-subjects analysis of variance (ANOVA). Locomotor activity and the time-course of cocaine responding were analyzed using two-way ANOVA, with repeated measures for both factors. Conditioned place preference was analyzed using Student’s *t*-test. Significant effects in the ANOVA were followed by the Newman-Keuls *post hoc* test. Values of *p* < 0.05 were considered statistically significant.

## Results

### Effect of cebranopadol on the escalation of cocaine self-administration

After 3 weeks of cocaine self-administration, the rats gradually escalated their cocaine consumption (one-way ANOVA, *F*_14,195_ = 8.498, *p* < 0.001). The Newman-Keuls *post hoc* test revealed significant escalation of cocaine self-administration that began from session 8 until session 15 compared with the first day of extended access (*p* < 0.01 for session 8; *p* < 0.001 for sessions 9–15; Fig. 1A). Treatment with cebranopadol significantly decreased operant responding for cocaine (*F*_2_,_13_ = 31.19, *p* < 0.001). The Newman-Keuls *post hoc* test revealed that cebranopadol reduced cocaine self-administration at both doses tested (*p* < 0.01, 25 μg/kg vs. 0.0 μg/kg; *p* < 0.001, 50 μg/kg vs. 0.0 μg/kg; Fig. 1B). Cebranopadol dose-dependently decreased the number of cocaine infusions (*p* < 0.01) to levels that were comparable to the pre-escalation baseline. When all of the data were included in a single statistical analysis (baseline pre-escalation, escalation doses of 0, 25, and 50 μg/kg), the one-way repeated-measures ANOVA revealed a significant effect of treatment (*F*_13,52_ = 4.080, *p* < 0.001). The Newman-Keuls *post hoc* test indicated no differences in intake between 50 μg/kg cebranopadol and the pre-escalation baseline. Inactive lever responses were very low and unaffected by cebranopadol (*F*_2,13_ = 0.089). Fig. 1C shows the time course of the cumulative number of cocaine infusions following cebranopadol treatment. Fig. 1D shows the time course of hourly infusions during the session, demonstrating that cebranopadol was effective over the entire 6 h self-administration session. The two-way repeated-measures ANOVA revealed significant effects of treatment (*F*_2_,_13_ = 31.19, *p* < 0.001) and time (*F*_5,65_ = 11.90, *p* < 0.001).

**Figure 1.**
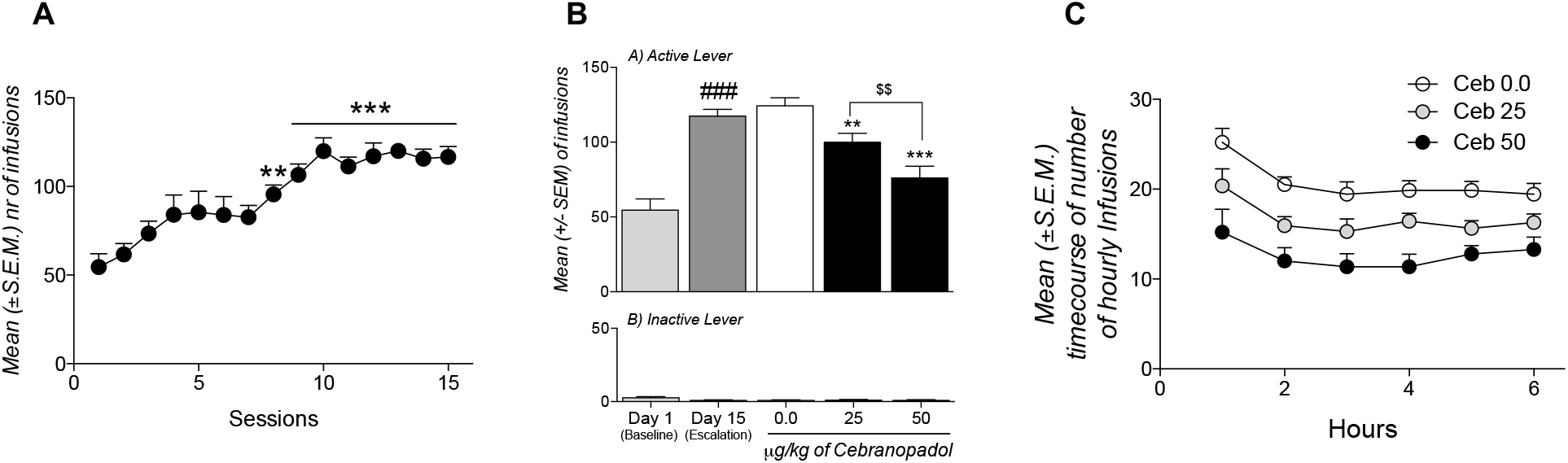
(**A**) Escalation of cocaine self-administration after 15 (6 h) sessions. \*\**p* < 0.01, \*\*\**p* < 0.001, significantly different from day 1. (**B**) Effect of cebranopadol on cocaine selfadministration. The data are expressed as the mean ± SEM number of responses on the active lever (A) and inactive lever (B). *n* = 14. \*\**p* < 0.01, \*\*\**p* < 0.001, significantly different from vehicle-treated rats (0.0 μg/kg). ^$^*p* < 0.05, 50 μg/kg vs. 25 μg/kg. (**C**) Hourly time course of infusions.

### Effect of cebranopadol on sweetened condensed milk self-administration

The data were analyzed at two different time points to compare the effects of cebranopadol on SCM and cocaine self-administration. We selected the 1 h time point to match the number of rewards between cocaine and SCM and the 6 h time point to match the length of the cocaine session. The one-way repeated-measures ANOVA revealed no effects of cebranopadol on SCM self-administration at 1 h (*F*_2,11_ = 0.63; Fig. 2A) or 6 h (*F*_2,11_ = 0.37; Fig. 2B).

**Figure 2.**
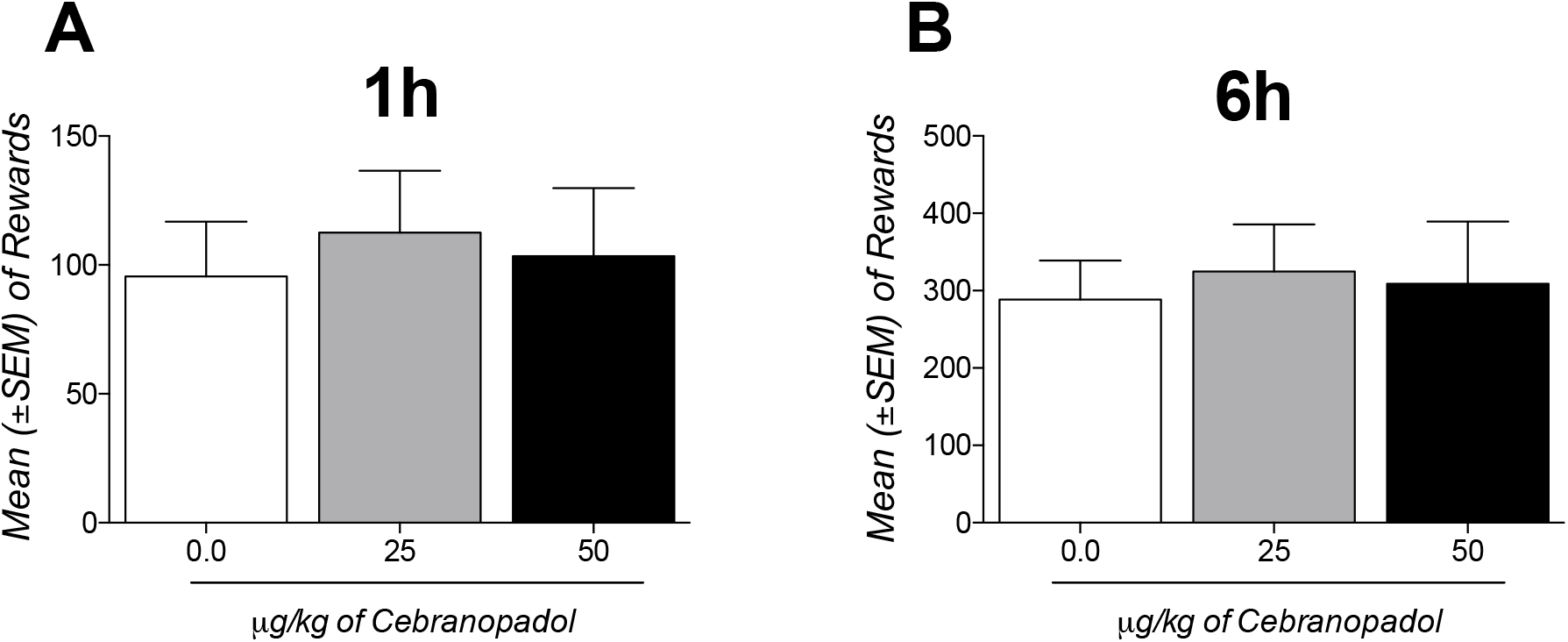
Effect of cebranopadol on SCM self-administration after 1 h (A) or 6 h (B). The data are expressed as the mean ± SEM number of rewards. *n* = 12.

### Cebranopadol-induced conditioned place preference and locomotor activity

We found no significant overall bias toward either of the compartments during the pretest, measured as the mean time spent in compartment A (812 ± 116 s) and compartment B (964 ± 117 s; *t*_8_ = 0.6557, *p* = 0.5304). The two-way repeated-measures ANOVA revealed no effects of cebranopadol on general locomotor activity during the conditioning sessions, with no effects of treatment (*F*_1,8_ = 0.008) or time (*F*_3,24_ = 2.8) and no treatment × time interaction (*F*_1,8_ = 0.33; Fig. 3A). However, cebranopadol induced conditioned place preference compared with the time spent in the drug-paired side during the pretest and preference test (*t*_8_ = 4.736,*p* < 0.01; Fig. 3B).

**Figure 3.**
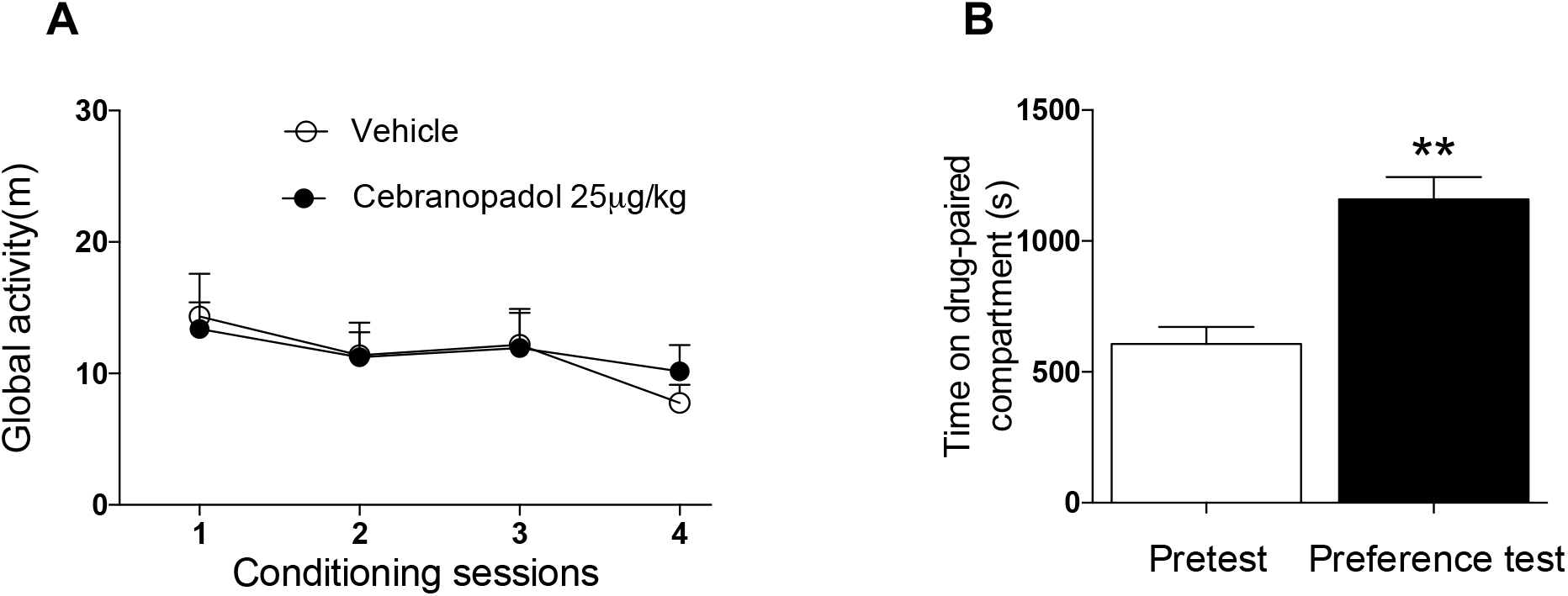
(**A**) Average total distance traveled (in meters) during the four sessions of CPP conditioning after oral injection of cebranopadol or vehicle. (**B**) Cebranopadol-induced conditioned place preference. The data are expressed as the time spent in the drug-paired chamber. *n* = 9.

### Effect of cebranopadol on the conditioned reinstatement of cocaine-seeking behavior

At the end of the cocaine self-administration phase, the average number of reinforced responses was 126 ± 6.2. During extinction, lever pressing decreased from 42.3 ± 8.3 during the first session to 10.2 ± 1.9 during the last extinction session (16 total sessions). As shown in Fig. 4A, reintroduction of the cocaine-discriminative cues (S^D^) but not neutral stimuli (S^N^) significantly reinstated extinguished cocaine-seeking behavior (*F*_2,9_ = 11.61, *p* < 0.001). Inactive lever presses were unaltered by cue presentation (*F*_2,9_ = 1.48; Fig. 4B). The one-way repeated-measures ANOVA revealed a significant effect of cebranopadol treatment on the conditioned reinstatement of cocaine-seeking behavior (*F*_2,9_ = 10.12, *p* < 0.001). The Newman-Keuls *post hoc* test indicated that oral treatment with 50 μg/kg cebranopadol 30 min before the reinstatement test significantly reduced cocaine-seeking behavior (*p* < 0.01, *vs*. 0.0 μg/kg; *p* < 0.05, *vs*. 25 μg/kg; Fig. 4A). Responses on the inactive lever remained negligible and unaltered by cebranopadol treatment (*F*_2,9_ = 0.82; Fig. 4B).

**Figure 4.**
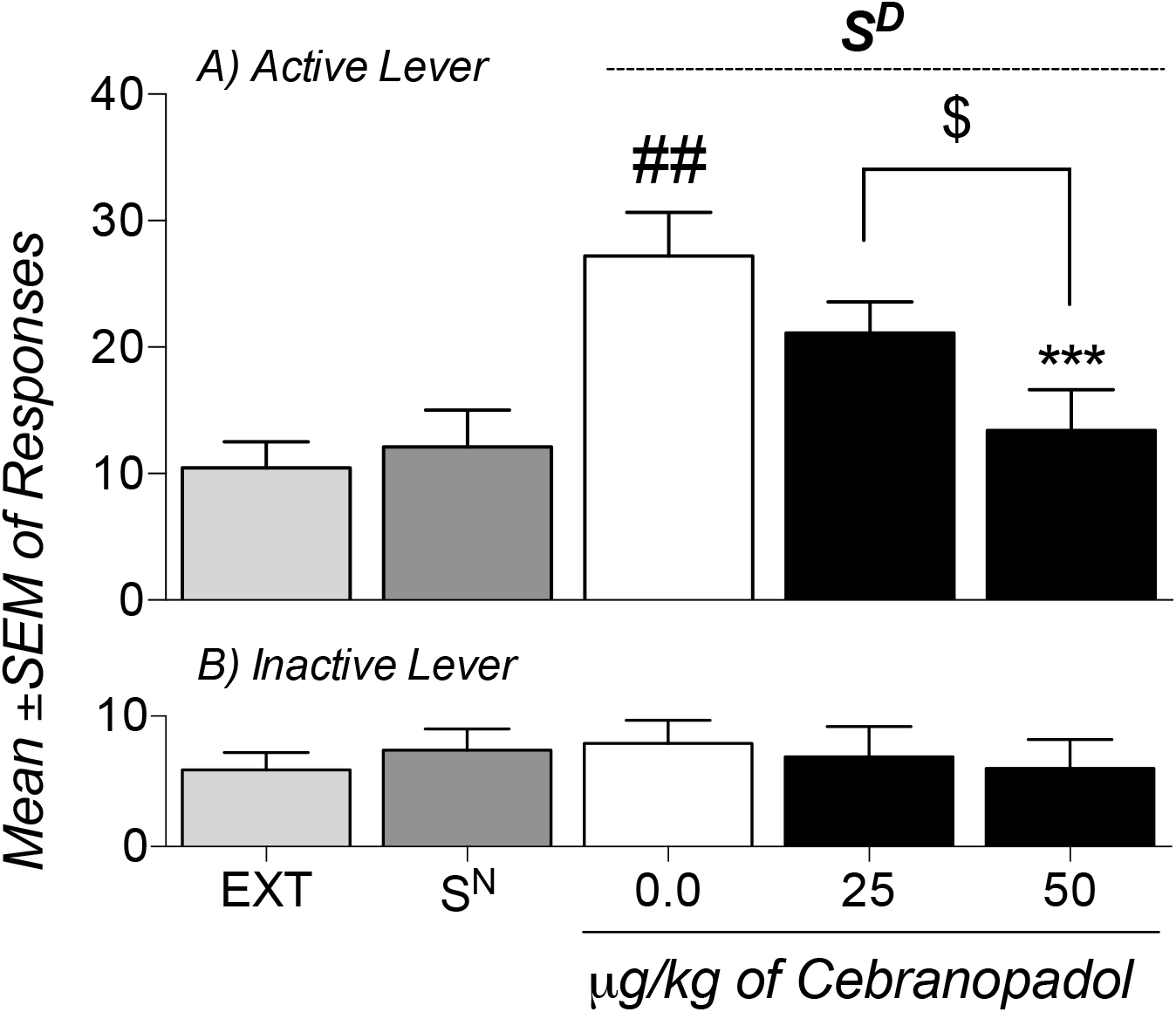
Effect of cebranopadol on the conditioned reinstatement of cocaine seeking. During the self-administration training phase, the rats learned to press the lever for cocaine in the presence of a contextual/discriminative stimulus (S^D^). During extinction (Ext), lever responding progressively decreased. Twenty-four hours after the last extinction session, all of the rats were presented with a neutral stimulus (S^N^), and the number of responses on the previous cocaine-paired lever was not reinstated. In the reinstatement test, responses to the cocaine-paired cues increased in the presence of the S^D^. Cebranopadol at the dose of 50 μg/kg significantly reduced relapse that was elicited by presentation of the S^D^. The data are expressed as the mean ± SEM number of responses on the cocaine active lever (A) and inactive lever (B). *n* = 10. ^≠≠≠^*p* < 0.001, significantly different from extinction; ^*\*\*\**^*p* < 0.01, significantly different from vehicle-treated rats (0.0 μg/kg); ^$^*p* < 0.05, significantly different from 25 μg/kg

## Discussion

The present study demonstrated that cebranopadol dose-dependently reduced cocaine self-administration in rats that had extended access to cocaine (Fig. 1B). Cebranopadol reduced cocaine intake during the entire 6 h session and did not affect inactive lever responding. Furthermore, it had no effect on the self-administration of a natural and highly palatable reinforcer (SCM). Finally, cebranopadol prevented the conditioned reinstatement of cocaine seeking, but it produced conditioned place preference.

The reduction of cocaine self-administration by cebranopadol is consistent with the effects of buprenorphine (a partial agonist at MORs and NOP receptors) on cocaine intake in rhesus monkeys (Mello et al., 1992) and rats (Sorge and Stewart, 2006a; Wee et al., 2012). However, buprenorphine and methadone are often associated with a decrease in locomotor activity (Marquez et al., 2007). In the present study, we did not observe any differences in locomotor activity after cebranopadol administration, suggesting that the decreases in cocaine self-administration and cocaine seeking in the reinstatement test were not attributable to nonspecific motor effects, and cebranopadol may have fewer undesirable side effects than buprenorphine and methadone.

The lack of effect of cebranopadol on SCM self-administration (compared with cocaine) did not likely result from different levels of responding for each reward because responses during the SCM self-administration were analyzed at two different time points to match both the number of rewards (1 h; Fig. 2A) and length of the cocaine and SCM self-administration sessions (6 h; Fig. 2B). Importantly, this selectivity of the effect of cebranopadol on cocaine self-administration was not reported with buprenorphine. Buprenorphine decreased the self-administration of both glucose and saccharin in rats (Carroll and Lac, 1992) and monkeys (Carroll et al., 1992). Therefore, the data suggest that cebranopadol has better selectivity in preventing behavior that is motivated by drugs of abuse over behavior that is directed toward a highly palatable food reward.

Nociceptin blocks morphine-, cocaine-, and alcohol-induced CPP (Ciccocioppo et al., 1999; Murphy et al., 1999; Ciccocioppo et al., 2002; Sakoori and Murphy, 2004), and it does not have any rewarding properties when administered alone (Devine et al., 1996). Additionally, NOP receptor agonists attenuate morphine-induced CPP (Shoblock et al., 2005). Intracerebroventricular nociceptin administration inhibited cocaine-induced CPP (Kotlinska et al., 2002). In contrast to these reports, recent data showed that genetic deletion of the nociceptin receptor in rats conferred resilience to abused psychoactive drugs (Kallupi et al., 2017), and NOP antagonism blocked both nicotine and alcohol self-administration in a co-administration paradigm (Cippitelli et al., 2016). One hypothesis is that these effects of NOP agonists are mediated by mechanisms that involve the downregulation or desensitization of NOP receptors (Ciccocioppo et al., 2014). Several groups have reported rapid and robust NOP receptor desensitization and internalization *in vitro* in response to nociceptin or NOP agonists (Dautzenberg et al., 2001; Corbani et al., 2004; Spampinato and Baiula, 2006).

Given the agonist properties of cebranopadol at both MORs and NOP receptors, we hypothesized that it would not induce CPP because NOP agonism would attenuate MOR-mediated rewarding effects. However, in the present study, cebranopadol induced CPP, suggesting that full agonist activity and equivalent potency at NOP receptors and MORs was insufficient to prevent the rewarding effects of MOR activation. Nevertheless, because of the biased CPP design, cebranopadol-induced CPP may reflect its rewarding properties and possible anxiolytic-like effects (Tzschentke, 1998). Similar findings were reported by Khroyan et al. (Khroyan et al., 2007), in which they characterized SR 16435, a compound that has a pharmacological profile that is similar to cebranopadol.

After the escalation of cocaine self-administration stabilizes, rats exhibit a very stable and reproducible pattern of behavior that is characterized by an initial loading phase, followed by the development of a regular response pattern and dose titration (Ahmed and Koob, 1998; Ahmed and Koob, 1999).When administered orally 30 min before the session, cebranopadol dose-dependently reduced the number of cocaine infusions during the loading phase (Fig. 1C). At the end of the loading phase, cebranopadol continued to dose-dependently reduce cocaine intake, demonstrating its long duration of action. One approach for treating cocaine addiction focuses on the use of compounds that have the same primary actions as the abused drug but a longer duration of action paired with lower intrinsic abuse potential and fewer toxic side effects (Kreek, 1997). We observed a long duration of action of cebranopadol (> 6 h), suggesting that cebranopadol may act similarly to buprenorphine and methadone as a slow-acting agonist/indirect agonist. Overall, cebranopadol may be a better candidate for substitution therapy for cocaine abuse because it appears to have fewer nonspecific effects on the motivation to seek and take natural rewards.

Environmental stimuli that are associated with cocaine use are major factors that can cause intense craving and precipitate cocaine relapse in humans. Stimuli that are predictive of the availability of cocaine reliably elicit strong craving and relapse of drug seeking, even after several months of abstinence (Weiss et al., 2001; Ciccocioppo et al., 2004; Martin-Fardon and Weiss, 2016). Cebranopadol significantly and dose-dependently reduced the cue-induced reinstatement of cocaine-seeking behavior, without affecting inactive responding. Cebranopadol decreased the reinstatement of cocaine seeking in the absence of cocaine under extinction conditions, demonstrating that it may decrease the motivation to seek and take cocaine, independent of changes in blood levels of cocaine. These results are consistent with the decrease in cocaine seeking that was induced by buprenorphine in the presence of drug-associated cues during extinction (Sorge et al., 2005).

One limitation of the present study was the lack of full characterization of the pharmacokinetics and pharmacodynamics of cebranopadol. We also did not identify shifts in the dose-response curve or specific receptors that mediate its preclinical efficacy. Follow-up studies are needed to fully characterize the reinforcing properties and possible abuse potential of cebranopadol, particularly considering that we found that cebranopadol produced conditioned place preference. However, although such characterization studies are important from a theoretical perspective to understand the precise mechanisms of action and facilitate medication development, cebranopadol has already been shown to be well tolerated in humans and is already being tested in several clinical trials for the treatment of pain.

In summary, the present study provides preclinical evidence of the efficacy of cebranopadol in reversing compulsive-like responding for cocaine and cue-induced reinstatement of cocaine seeking. Cebranopadol may be a new therapeutic option for the prevention of cocaine abuse and relapse.

## Acknowledgements

The authors thank Michael Arends for proofreading the manuscript.

## Author Contributions

Participated in research design: de Guglielmo, Martin-Fardon, George. Conducted experiments: de Guglielmo, Matzeu, Kononoff, Mattioni. Performed data analysis: de Guglielmo. Wrote or contributed to writing the manuscript: de Guglielmo, Martin-Fardon, George.

## Footnotes

This study was supported by National Institutes of Health [Grant AA006420], [Grant AA020608], [Grant AA022977] (OG), [Grant DA033344], and [Grant AA024146] (RMF) and the Sigrid Juselius Foundation (JK). The authors declare no conflict of interest.

